## Supplementary Figures and Text for "Echolocation reverses information flow in a cortical vocalization network"

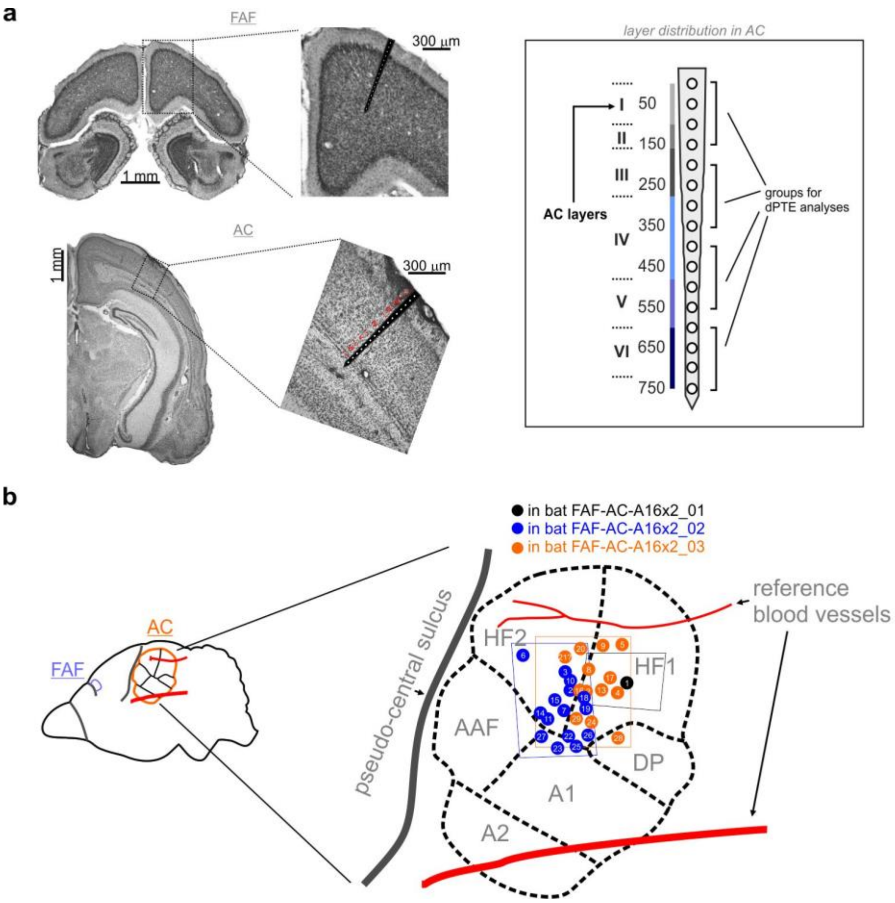


**Fig. S1. Location of recordings in *C. perspicillta*'s auditory cortex**. (**a**) Coronal sections of the frontal auditory field (FAF; top) and the auditory cortex (AC; bottom) of *C. perspicillata*. In the zoomed-in insets for each location, an in-scale schematic of the electrode positioning in shown. For the case of the auditory cortex, layer borders are demarcated (see also rightmost inset of the panel); for the case of the FAF, layer borders were difficult to determine and are therefore not shown. Note, in the rightmost inset of the panel, the grouping for the dPTE analyses relative to electrode depth and AC laminar distribution. (**b**) Laminar recordings in the AC were made mostly in the high frequency fields. In the panel, the approximate position of each recording is illustrated. Each circle corresponds to a single penetration, each being of three possible colours depending on the animal where experiments were performed. Areas delimited by orange, black or blue dashed lines (each colour corresponding to a specific animal) depict the approximate extent of the trepanations.

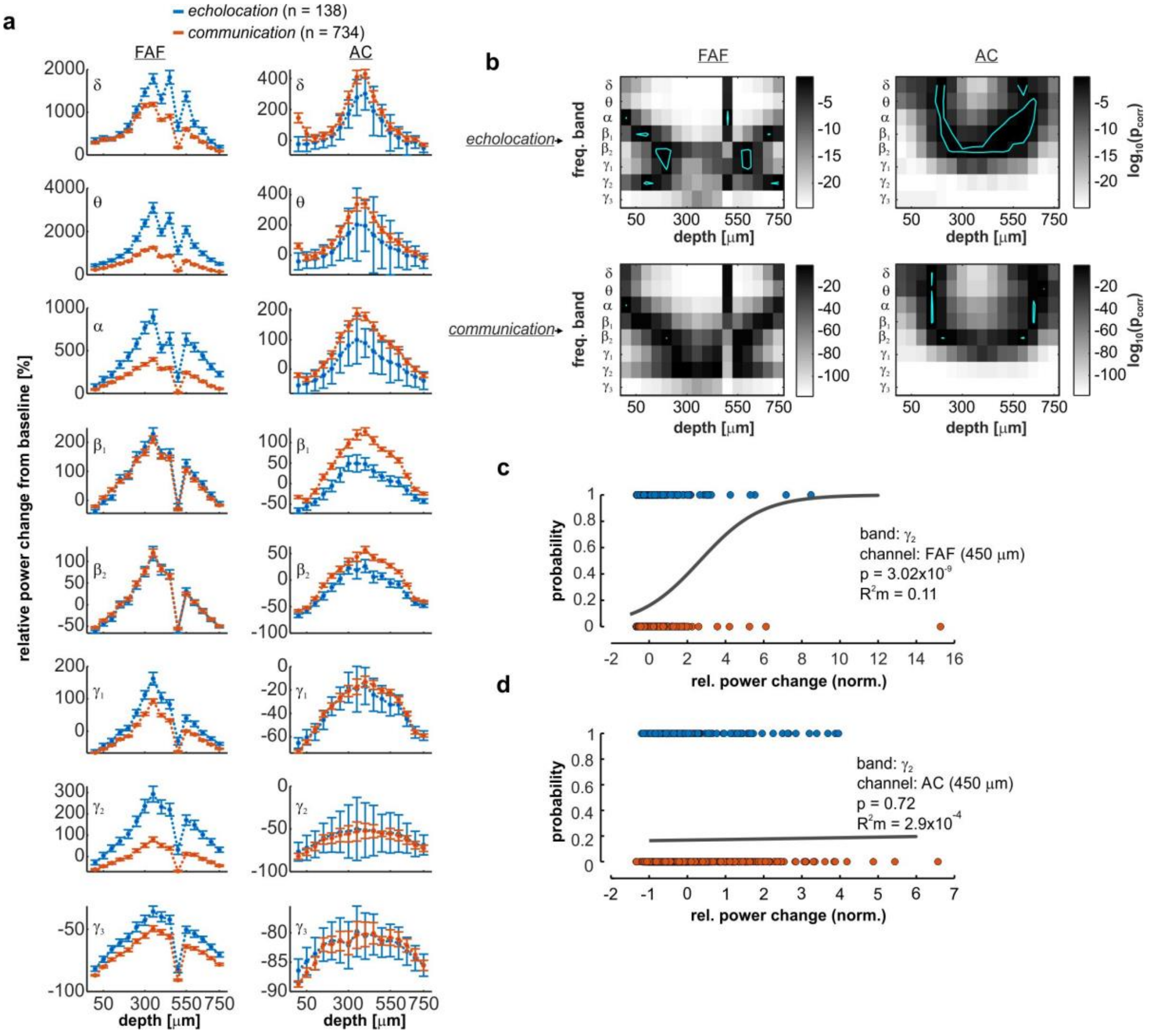


**Fig. S2. Pre-vocal power changes associated with the production of echolocation and communication calls**. (a) Percentage pre-vocal power change across LFP frequency bands ($\delta$, 1-4 Hz; $\theta$, 4-8 Hz; $\alpha$, 8-12 Hz; $\beta_1$, 12-20 Hz; $\beta_2$, 20-30 Hz; $\gamma_1$, 30-60 Hz; $\gamma_2$, 60-120 Hz; $\gamma_3$, 120-200 Hz), relative to a no-voc baseline, across all cortical depths in FAF (left) and AC (right). Pre-vocal power change values related to echolocation utterances (n = 138) are depicted in blue; those related to communication utterances (n = 734) are depicted in orange. Data shown as mean ± sem. (**b**) Significance matrices depicting p-values statistical tests to determine whether changes shown in panel a were significant (i.e. significantly different than 0% change for each channel and frequency band; FDR-corrected Wilcoxon signed rank tests). The colour scale in the figures indicates the log10 of the corrected p-values (significance when $p_{corr} < 0.05$). (**c**) Example GLM fitted with pre-vocal power change data from an FAF channel located at 450 mm from the cortical surface, in the $g_2$-band. Power changes in this band significantly predicted ensuing call type on a trial-by-trial basis ($p = 3.02x10^{-9}$), with moderate effect size $R^2m = 0.11$. (**d**) Example GLM fitted with pre-vocal power change data from the AC, same electrode depth as in c, and also in the $g_2$ frequency band. Relative power changes in this frequency band and brain region did not significantly predict ensuing vocal type (p = 0.72).

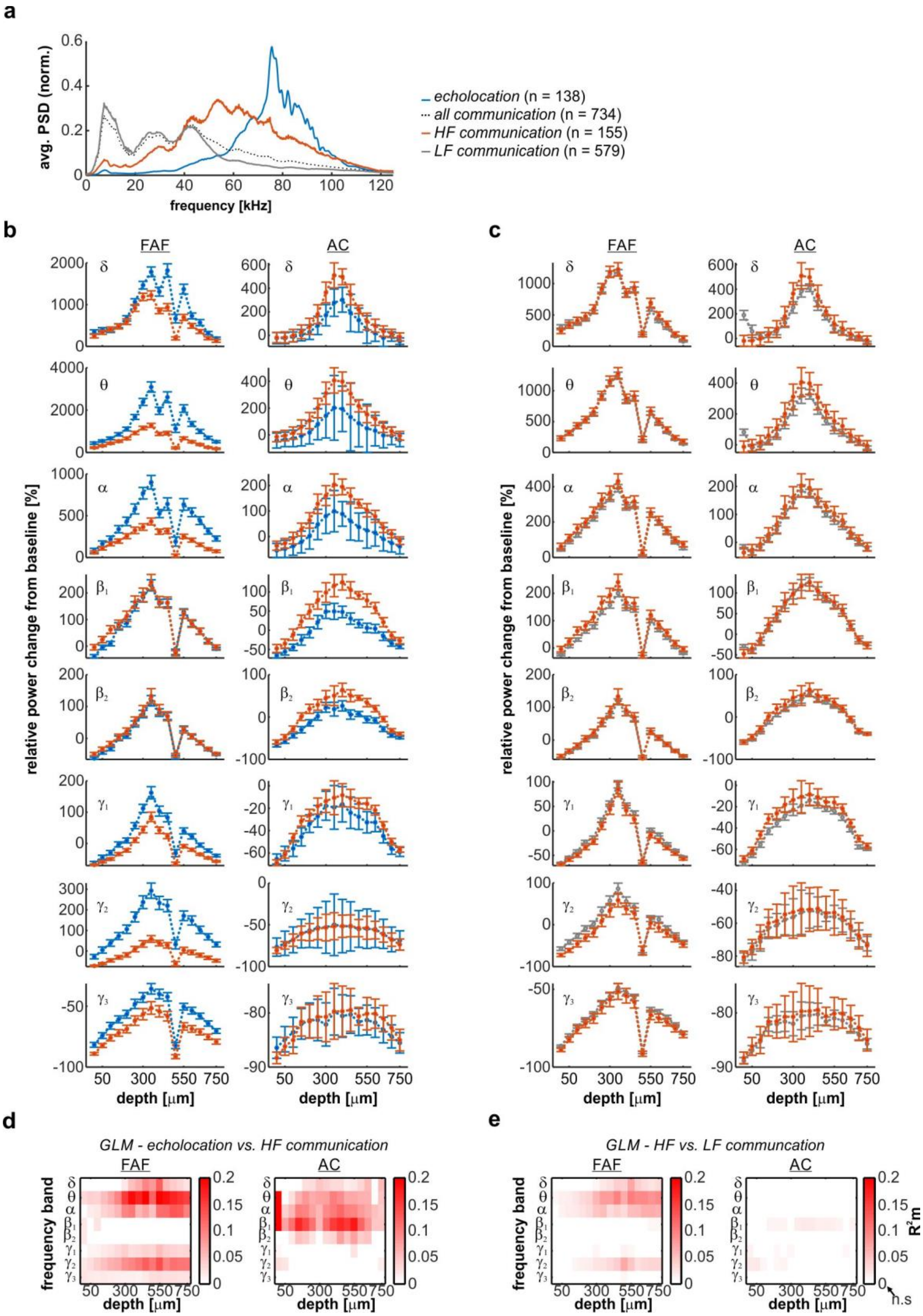
a
avg. PSD (norm.)
frequency [kHz]
echolocation (n = 138)
all communication (n = 734)
HF communication (n = 155)
LF communication (n = 579)
b
c
FAF
AC
relative power change from baseline [%]
depth [μm]
d
GLM - echolocation vs. HF communication
e
GLM - HF vs. LF communcation
frequency band
$R^2m$
n.s

**Fig. S3. Pre-vocal LFP power distinguishes between ensuing HF-communication and echolocation calls**. (**a**) Normalized average power spectral density (PSD) of echolocation (blue), HF-communication (orange), LF-communication (gray), and all communication calls (black, dashed). (**b**) Percentage pre-vocal power change across LFP frequency bands ($\delta$, 1-4 Hz; $\theta$, 4-8 Hz; $\alpha$, 8-12 Hz; $\beta_1$, 12-20 Hz; $\beta_2$, 20-30 Hz; $\gamma_1$, 30-60 Hz; $\gamma_2$, 60-120 Hz; $\gamma_3$, 120-200 Hz), relative to a no-voc baseline, across all cortical depths in FAF (left) and AC (right). Pre-vocal power change values related to echolocation utterances (n = 138) are depicted in blue; those related to HF-communication utterances (n = 155) are depicted in orange. Data shown as mean ± sem. Pre-vocal LFP power differences between HF-communication and echolocation conditions resemble the patterns shown in **Fig. 1**. (**c**) Same as in **b**, but the data shown corresponds to the pre-vocal power change across LFP bands associated to the vocalization of HF-communication (orange) and LF-communication (gray, n = 579) calls. Note that differences across HF- and LF-communication calls are barely perceptible. Interestingly, pre-vocal LF-communication calls appeared higher than for HF-communication calls, strengthening the notion that the utterance of high frequency sounds does not necessarily imply higher power in pre-vocal LFPs. (**d**) Pre-vocal power change in frontal and auditory regions predict whether animals vocalize HF-communication or echolocation calls. Effect size ($R^2_m$) of GLMs considering all frequency bands and channels, both in frontal and auditory cortices. Effect sizes were considered small when $R^2_m < 0.1$, and medium for $R^2_m >= 0.1$. For illustrative purposes, effect size values from non-significant models were set to 0. (**e**) Same as in **d**, but depicting GLM outcomes for classifying across the HF- and LF-communication conditions. Models in FAF and predict ensuing vocal type poorly ($R^2_m <= 0.1$, i.e. small effect sizes).

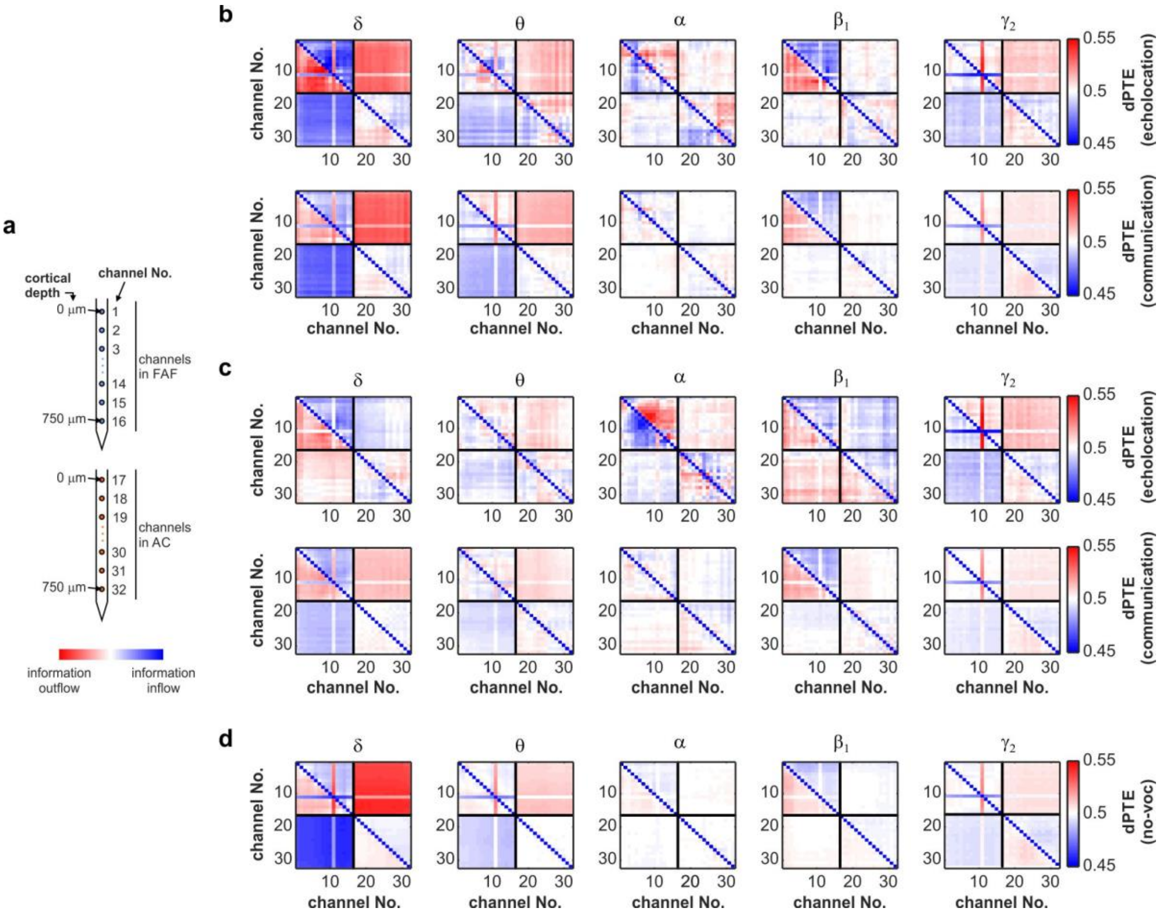


**Fig. S4. Average dPTE matrices during pre-vocal, post-vocal, and no-voc periods.** (**a**) Schematic representation of channel depth and cortical region associated with channel numbers in the panel. (**b**) Mean pre-vocal directed phase transfer entropy (dPTE) across LFP frequency bands ($\delta$, $\theta$, $\alpha$, $\beta_1$, $\gamma_2$) and conditions (echolocation utterance, top; communication utterance, bottom); 500 repetitions each). (**c**) Same as in **b**, with dPTE data corresponding to post-vocal periods. (**d**) Similar to **b** and **c**, illustrating average dPTE matrices corresponding to no-voc periods. Each matrix in the figure (i.e. panels **b-d**) illustrates the average dPTE across 500 repetitions calculated using 50 trials corresponding to echolocation, communication (both pre- and post-vocal), or no-voc related LFP segments. A cell (*i, j*) in a matrix shows the average dPTE value related to the information flow between channels *i* and *j*, which occurs in the $i \rightarrow j$ direction for dPTE values > 0.5 (red colours), and in the $j \rightarrow i$ direction for dPTE values < 0.5 (blue colours).

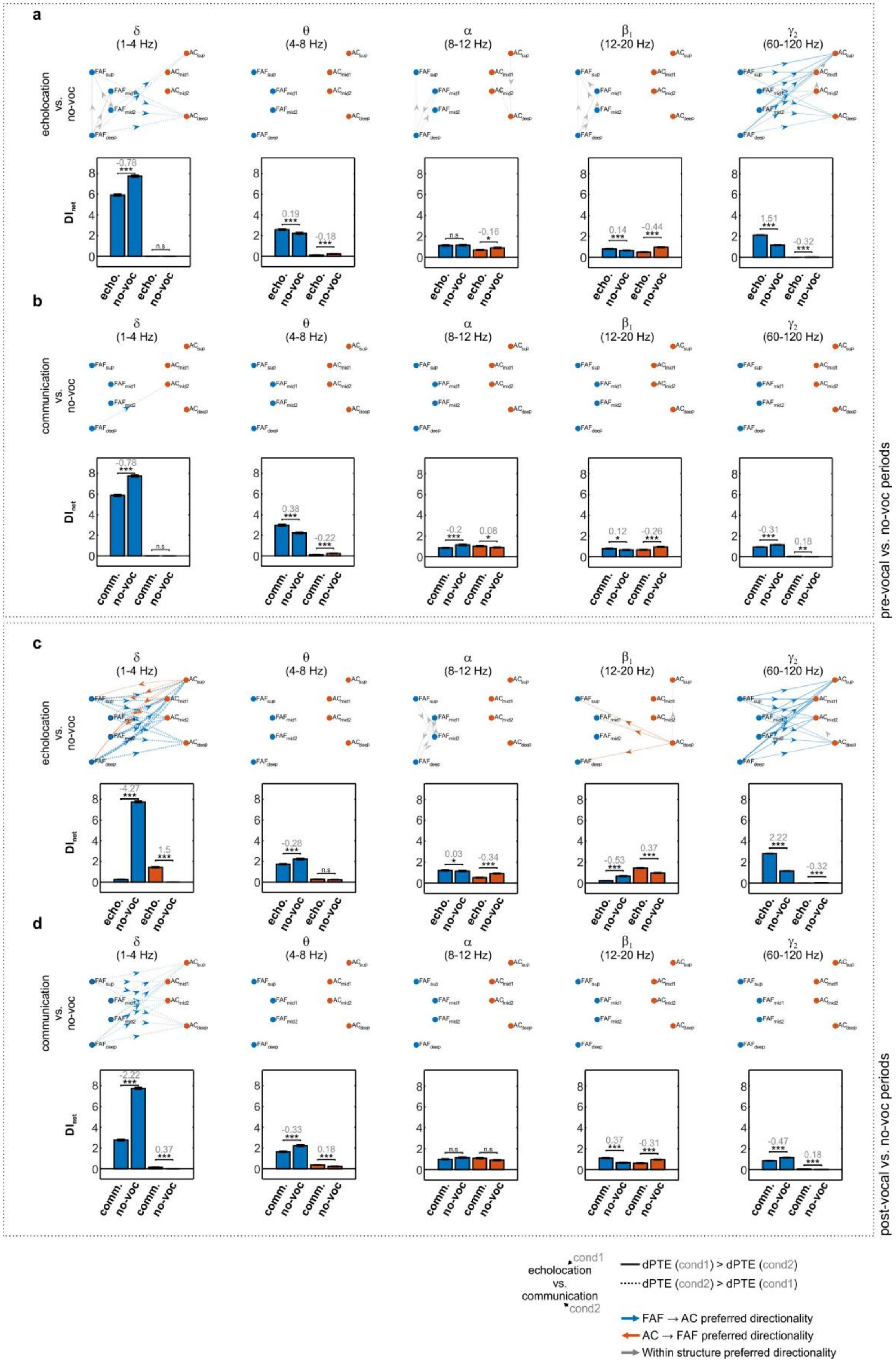
a
δ (1-4 Hz)
θ (4-8 Hz)
α (8-12 Hz)
β1 (12-20 Hz)
γ2 (60-120 Hz)
echolocation vs. no-voc
FAFsup FAFmid1 FAFmid2 FAFdeep
ACsup ACmid1 ACmid2 ACdeep
DInet
-0.78 *** n.s
0.19 *** -0.18 ***
n.s -0.16 *
0.14 *** -0.44 ***
1.51 *** -0.32 ***
echo. no-voc echo. no-voc
b
communication vs. no-voc
-0.78 *** n.s
0.38 *** -0.22 ***
-0.2 *** 0.08 *
0.12 * -0.26 ***
-0.31 *** 0.18 **
comm. no-voc comm. no-voc
pre-vocal vs. no-voc periods
c
echolocation vs. no-voc
-4.27 *** 1.5 ***
-0.28 *** n.s
0.03 * -0.34 ***
-0.53 *** 0.37 ***
2.22 *** -0.32 ***
echo. no-voc echo. no-voc
d
communication vs. no-voc
-2.22 *** 0.37 ***
-0.33 *** 0.18 ***
n.s n.s
0.37 *** -0.31 ***
-0.47 *** 0.18 ***
comm. no-voc comm. no-voc
post-vocal vs. no-voc periods
cond1
echolocation
vs.
communication
cond2
dPTE (cond1) > dPTE (cond2)
dPTE (cond2) > dPTE (cond1)
FAF → AC preferred directionality
AC → FAF preferred directionality
Within structure preferred directionality

**Fig. S5. Information flow differences in the FAF-AC circuit between vocalization and no-voc conditions.** (**a**, **b**) (*Top*) Graphs illustrating the differences between pre-vocal directionality and no-voc periods (**a**, echolocation vs. no-voc; **b**, communication vs. no-voc), across frequency bands. Edges were shown if three conditions were met: (i) the differences were significant (FDR-corrected Wilcoxon rank sum tests, $p_{corr} < 0.05$), (ii) the effect size was large ($|d| > 0.8$), and (iii) edges had already shown significant directionality (see edges in **Fig. 2**). Edge thickness is weighted according to the effect size of the comparison. Continuous lines indicate dPTEs for the first condition considered (see labels) higher than dPTEs for the second condition. Dashed lines indicate the opposite. (*Bottom*) Net information outflow ($DI_{net}$) from FAF (blue bars) and AC (orange bars), in the two conditions considered (**a**, echolocation vs. no-voc; **b**, communication vs. no-voc). Significant differences across conditions are marked with stars (FDR-corrected Wilcoxon rank sum tests; * $p_{corr} < 0.05$, ** $p_{corr} < 0.01$, *** $p_{corr} < 0.001$, n.s.: not significant; n = 500). Grey numbers in the panels indicate effect sizes (d; not shown for non-significant differences). Values were considered independently of whether there was previous significant directionality in any of the two conditions. Data shown as mean ± sem. (**c**, **d**) Same as **a**-**b**, but comparisons were made between post-vocal and no-voc periods (i.e. **c**, echolocation vs. no-voc; **d**, communication vs. no-voc).

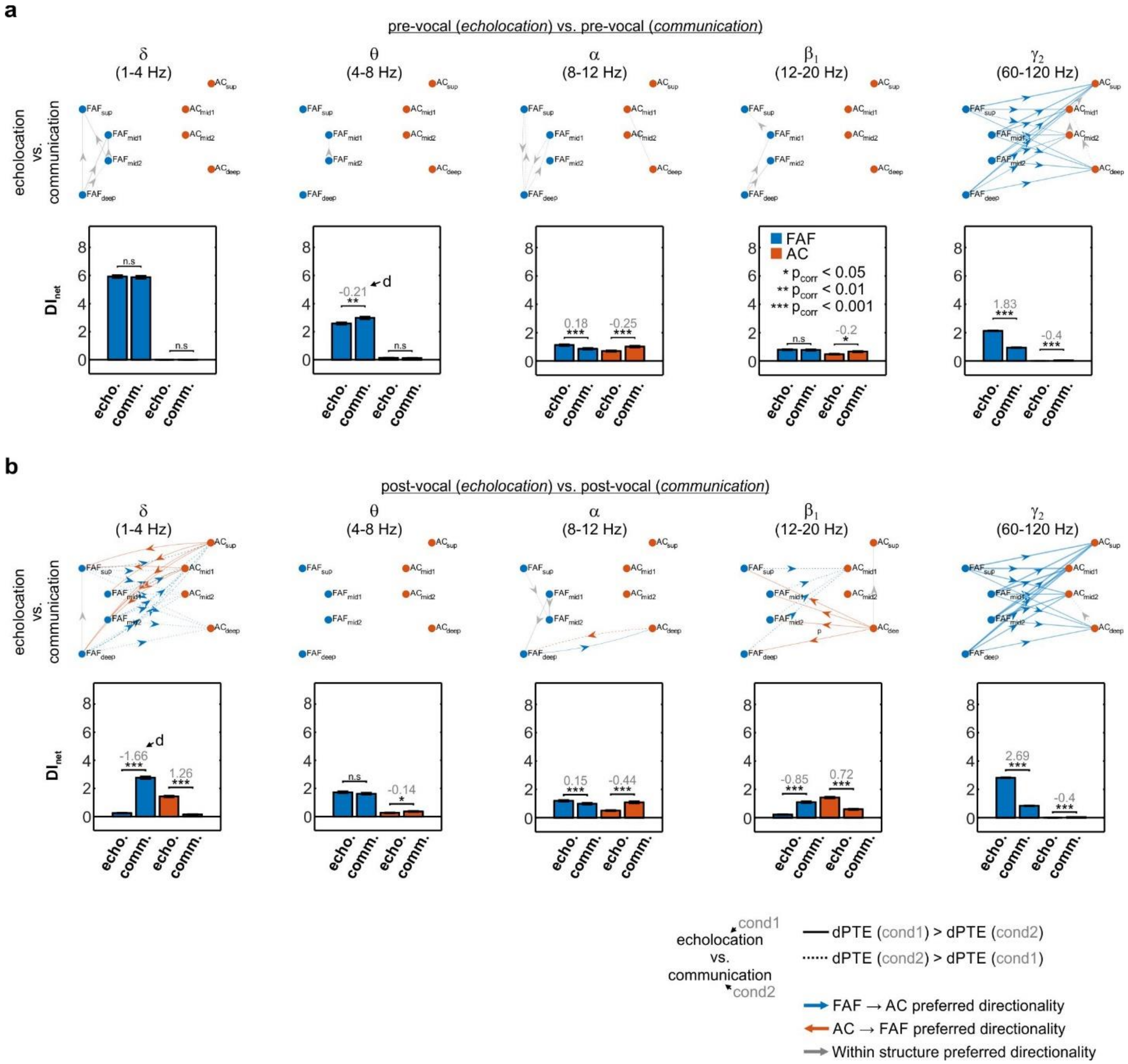


**Fig. S6. dPTE differences across vocalization conditions during pre-vocal and post-vocal periods.** (**a**) Differences in pre-vocal directionality of information flow between vocalization conditions (echolocation vs. communication), across frequency bands. Conventions for this figure are the same as for **Fig. S5**. (**b**) Same as **a**, but data shown correspond to comparisons between post-vocal periods related to echolocation and communication call production.

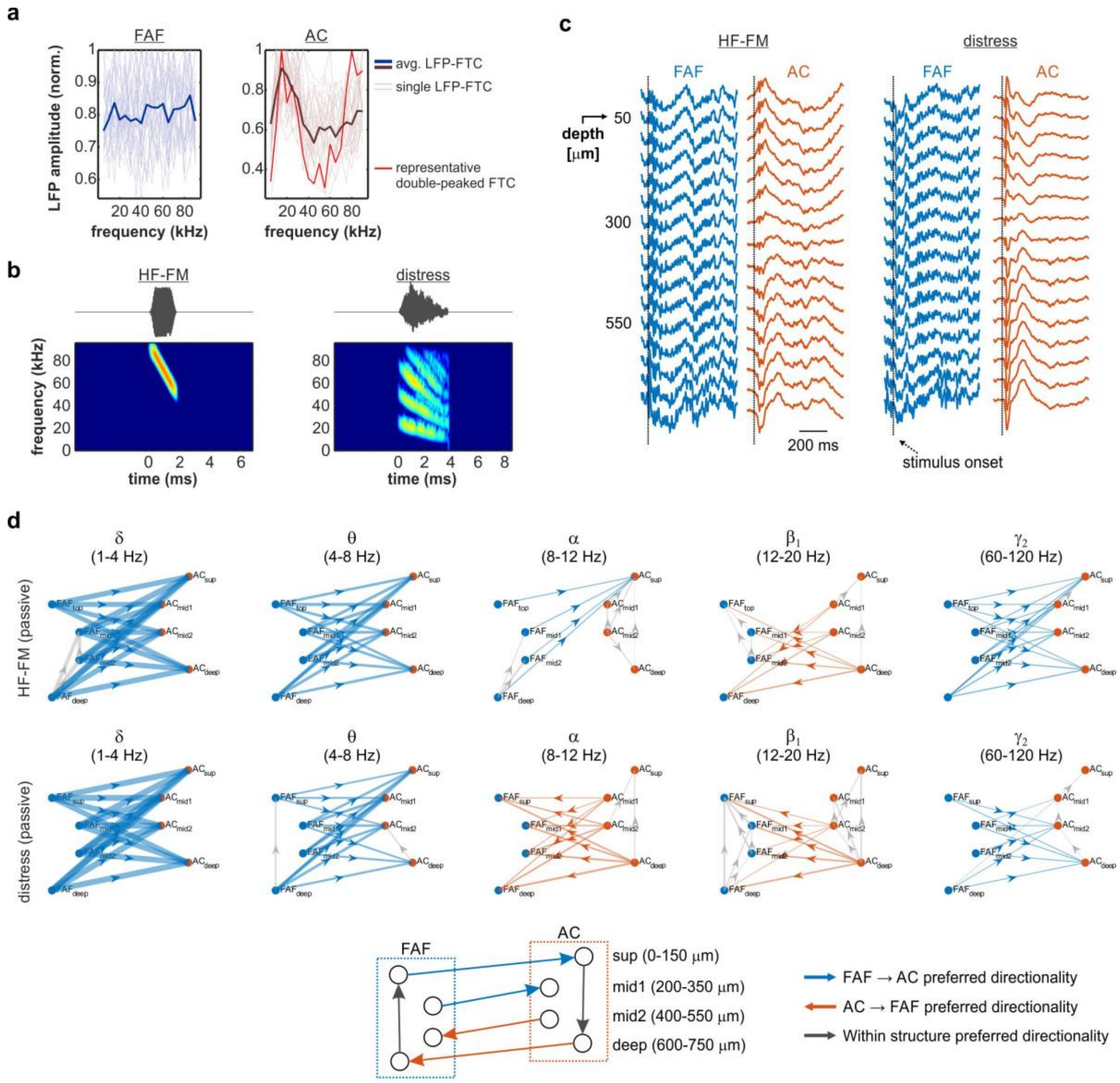


**Fig. S7. Directed connectivity in the FAF-AC circuit during passive listening.** (**a**) Frequency tuning curves (FTC) at 75 dB SPL for LFPs recorded in FAF and AC (at a representative depth of 350 μm). (**b**) Oscillograms (top) and spectrograms (bottom) of the sounds used for acoustic stimulation (HF-FM: high-frequency frequency-modulated). (**c**) Representative responses from FAF and AC (averaged across trials for one penetration pair) to the sounds presented in the passive listening experiments (HF-FM and distress). (**d**) Graph visualization of directed connectivity in the FAF-AC circuit during passive listening (period of 500 ms after sound onset; top: responses to HF-FM sweep, bottom: responses to distress syllables). Graph visualization follows the conventions described for **Fig. 2**. (See Supplementary Results).

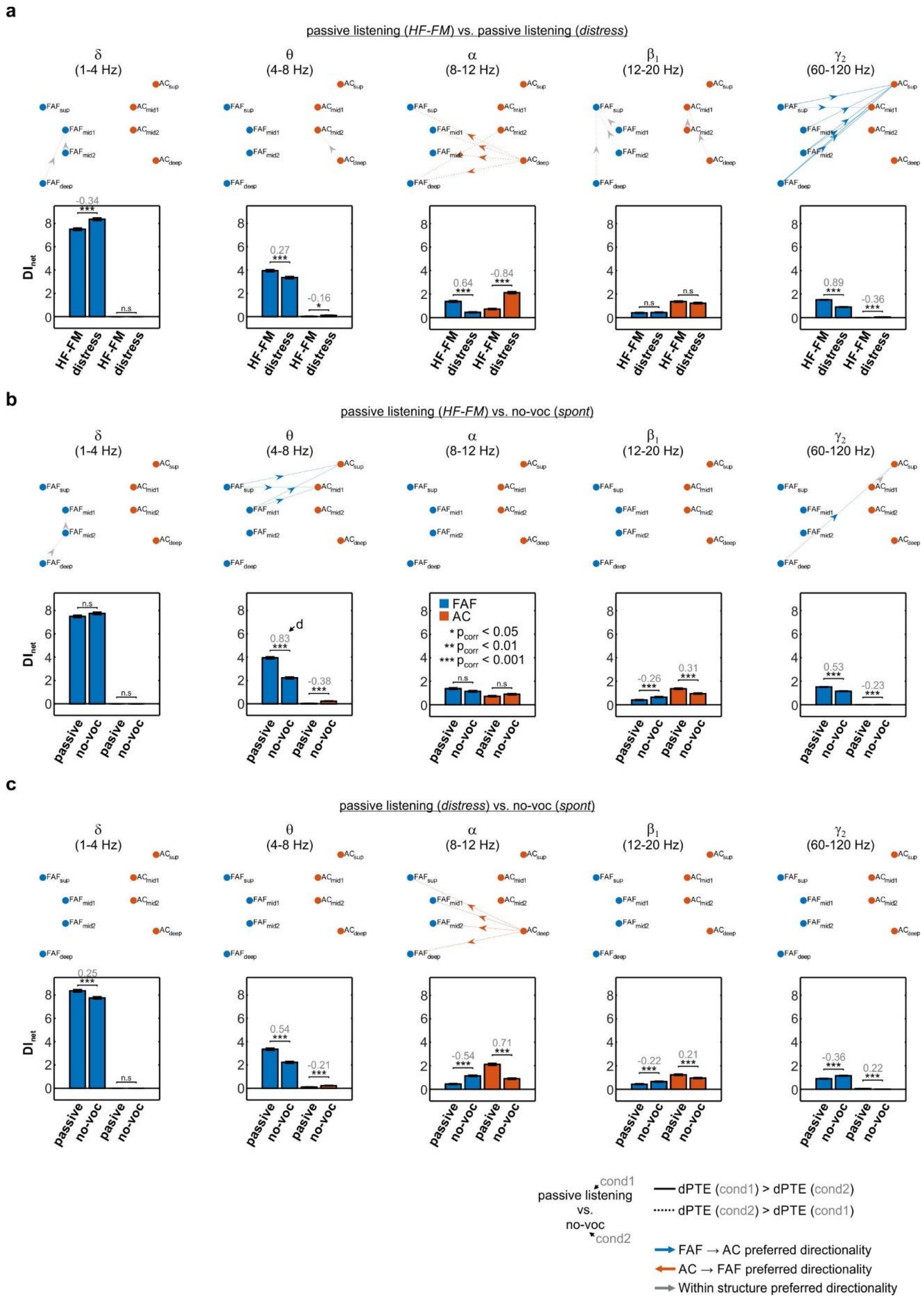


**Fig. S8. Information flow differences in the FAF-AC circuit between passive listening and no-voc conditions.** (**a**) Graph-based comparison of preferred directionality of information flow across passive listening conditions (i.e. responses to HF-FM sounds vs. responses to distress sounds). Conventions in the panel are the same as those described for **Fig. S5a**. (**b**) Same as in **a**, but data show comparisons between passive listening of HF-FM sounds vs. spontaneous activity (no-voc periods). (**c**) Same as in **b**, but data correspond to comparisons between passive listening of distress syllables vs. no-voc periods. (See Supplementary Results).

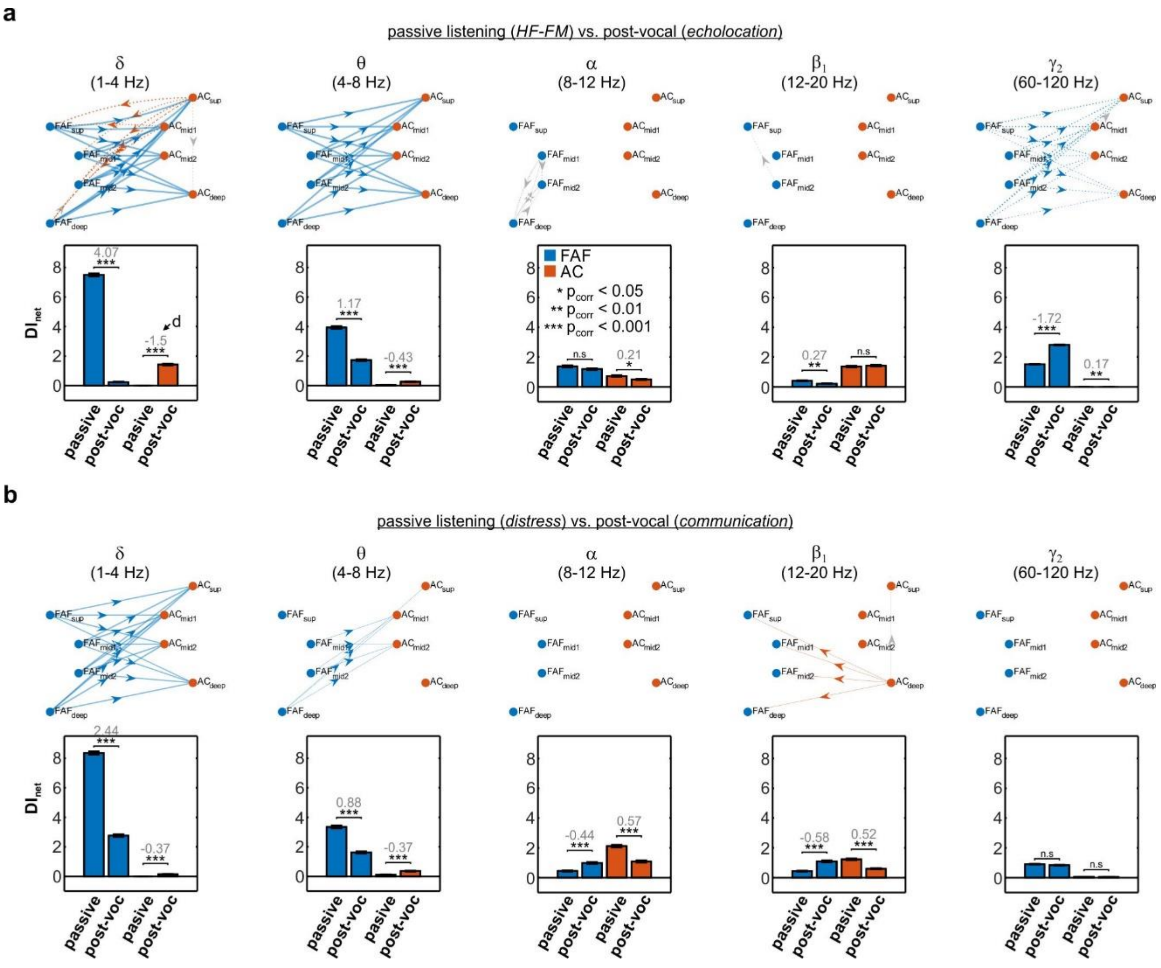


**Fig. S9. Directed connectivity differences across vocalizing and passively listening conditions.** (**a**) (*Top*) Graphs illustrating the differences in dPTE across passive listening and vocalization conditions (HF-FM sweep vs. active echolocation), across frequency bands. Edges shown follow the guidelines of those in **Fig. S5** (note that the basis for significant edges stem from **Fig. S6**). Continuous lines indicate dPTEs for passive listening of HF-FM sounds (first condition) higher than dPTEs for echolocation production (second condition). Dashed lines indicate the opposite. (*Bottom*) Net information outflow ($DI_{net}$) from FAF (blue bars) and AC (orange bars), in the two conditions considered. Data are shown following the conventions described in **Fig. S5**. (**b**) Same as in a, but comparing between passive listening of distress syllables vs communication production. (See Supplementary Results).

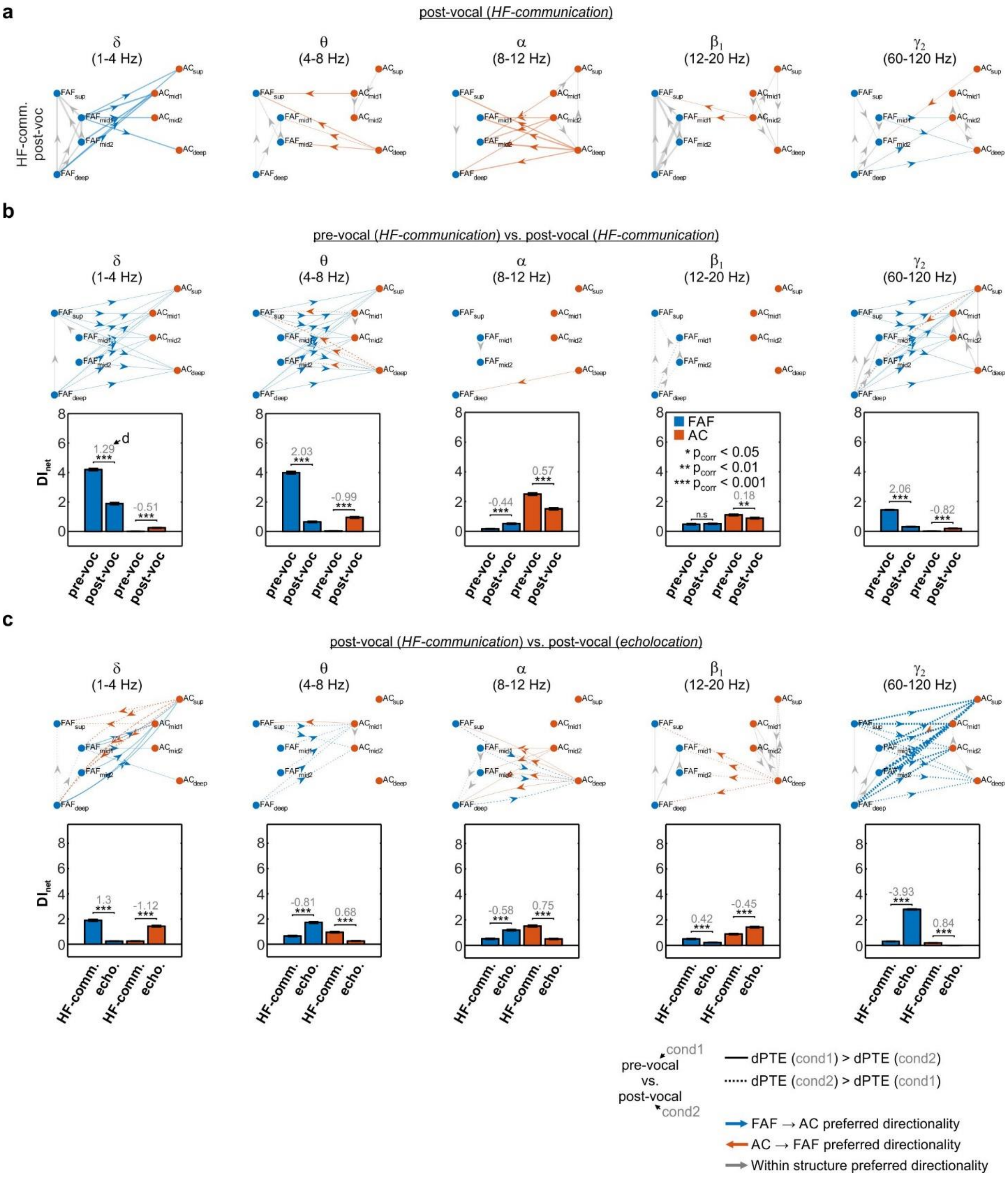

**Fig. S10. Information flow patterns during HF-communication production differ from those during echolocation production.** (**a**) Graph visualization of directed connectivity in the FAF-AC circuit for post-vocal periods associated to the production of HF-communication calls. Edges were shown according to the conventions described in **Fig. S5**. (**b**) (*Top*) Graphs illustrating the differences in dPTE across pre-vocal and post-vocal periods related to the production of HF-communication calls. Edge comparisons were made as described for **Fig. S5**. (*Bottom*) Net information outflow ($DI_{net}$) from FAF (blue bars) and AC (orange bars), in the two conditions considered. Data are shown as described for **Fig. 3**. (**c**) Same as in **b**, but comparisons were made between post-vocal periods related to the utterance of HF-communication (condition 1) and echolocation (condition 2) vocalizations. (See Supplementary Results).

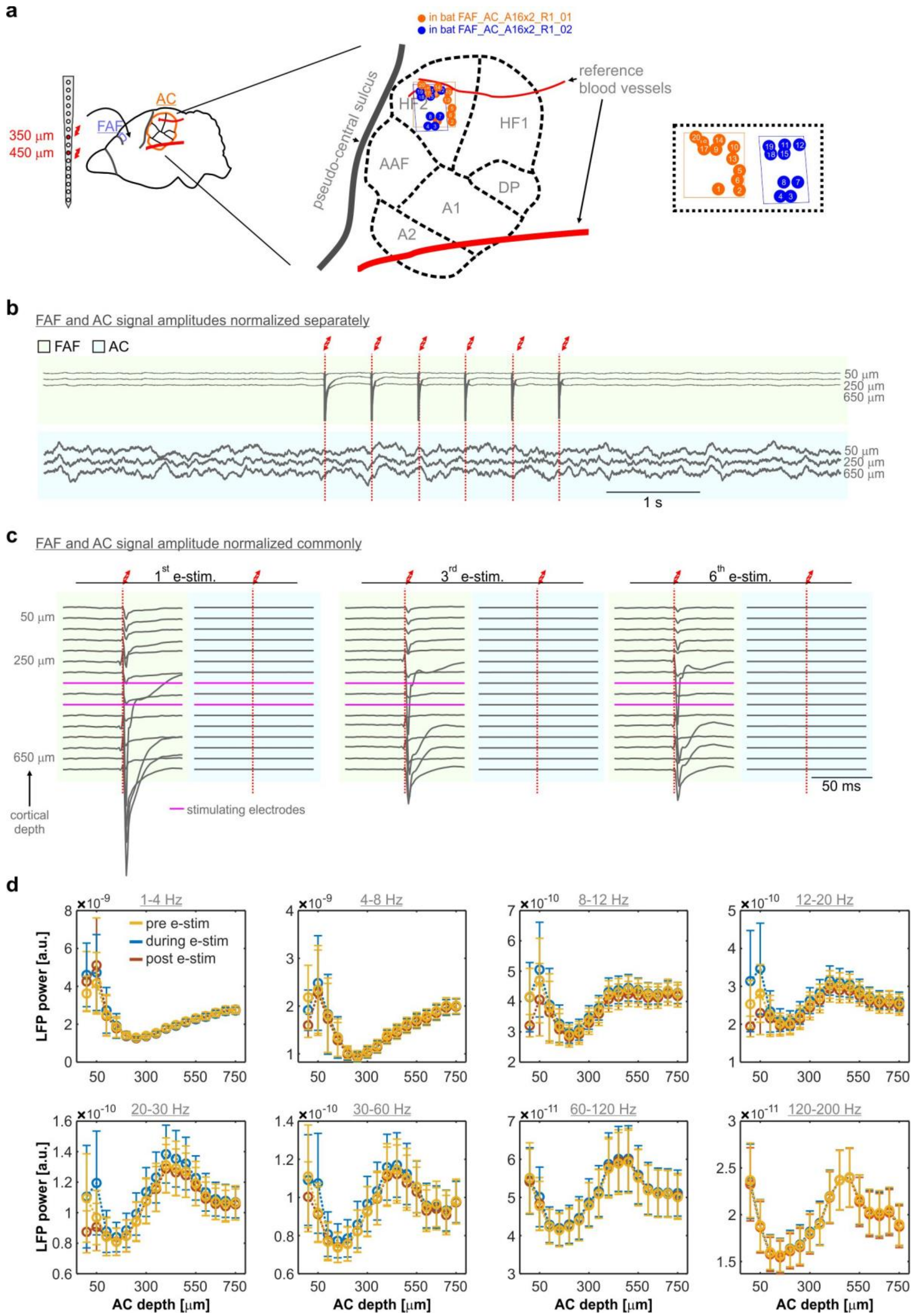
a
in bat FAF_AC_A16x2_R1_01
in bat FAF_AC_A16x2_R1_02
AC
FAF
350 μm
450 μm
pseudo-central sulcus
reference blood vessels
HF2
HF1
AAF
DP
A1
A2
b
FAF and AC signal amplitudes normalized separately
FAF
AC
50 μm
250 μm
650 μm
1 s
c
FAF and AC signal amplitude normalized commonly
1st e-stim.
3rd e-stim.
6th e-stim.
50 μm
250 μm
650 μm
cortical depth
stimulating electrodes
50 ms
d
1-4 Hz
4-8 Hz
8-12 Hz
12-20 Hz
20-30 Hz
30-60 Hz
60-120 Hz
120-200 Hz
pre e-stim
during e-stim
post e-stim
LFP power [a.u.]
AC depth [μm]

**Fig. S11. Electrical stimulation in FAF does not lead to consistent changes in concomitant auditory cortical LFPs.** (**a**) Schematic representation of electrical stimulation in FAF and recording locations in AC. The illustration of auditory cortical locations follows the same guidelines than those in **Fig. S1a** in this document. A zoom-in into the recording locations, separated by animals, is given in the rectangle to the rightmost part of the panel. (**b**) Grand average (calculated over a total of 1000 trials (i.e. 50 trials in 20 penetratios)) of LFPS around the times of electrical stimulation in FAF. Signals in this panel were normalized within structure: frontal and auditory cortical LFP amplitudes are comparable within each region, but not across. For illustrative purposes, simultaneously recorded FAF and AC field potentials are shown at three representative depths per structure (50, 250, 650 μm). Note that, in the FAF, the electrical artefact produced by the electrical stimulation is dominant. In the AC, however, no effects from the electrical stimulation are evident. (**c**) Grand average LFPs from AC and FAF (see **b**), but signal amplitude was normalized commonly across structures. That is, amplitudes in FAF and AC are comparable to one another. All recording depths are depicted, but only LFPs around the times of the first, third, and sixth electrical pulse (in a 6-pulse train) are shown. Note that electrical artefacts in the FAF are predominant when an electrical pulse is delivered, but that the amplitude of the AC average is considerably lower in comparison. (**d**) Comparisons of LFP power in AC before, during, and after electrical stimulation of the FAF across frequency bands (indicated in gray, above each sub-panel). The LFP power was obtained per penetration, and calculated as the average power across trials (n = 50). The time windows were as follows: for pre electrical stimulation (pre e-estim) segments, -2700 to -200 ms relative to first pulse onset; for segments during electrical stimulation (during e-stim), 0 to 2500 ms relative to first pulse onset; for segments post electrical stimulation (post e-stim), 2700 to 5200 ms relative first pulse onset. In other words, all segments were 2.5 seconds long, the same duration as the electrical pulse train. Band power was obtained by integrating LFP power between the given frequencies of a band (same procedure used for the data shown in **Fig. 1**). We observed no significant differences between the LFP power of pre-, during, and post- e-stim segments in AC, regardless of the frequency band considered. These results (panels **b**-**d**) indicate that the electrical stimulation of the FAF did not significantly alter auditory cortical LFPs by means of entrainment to the electrical stimulation train, or other passive propagation of electrical artefacts.

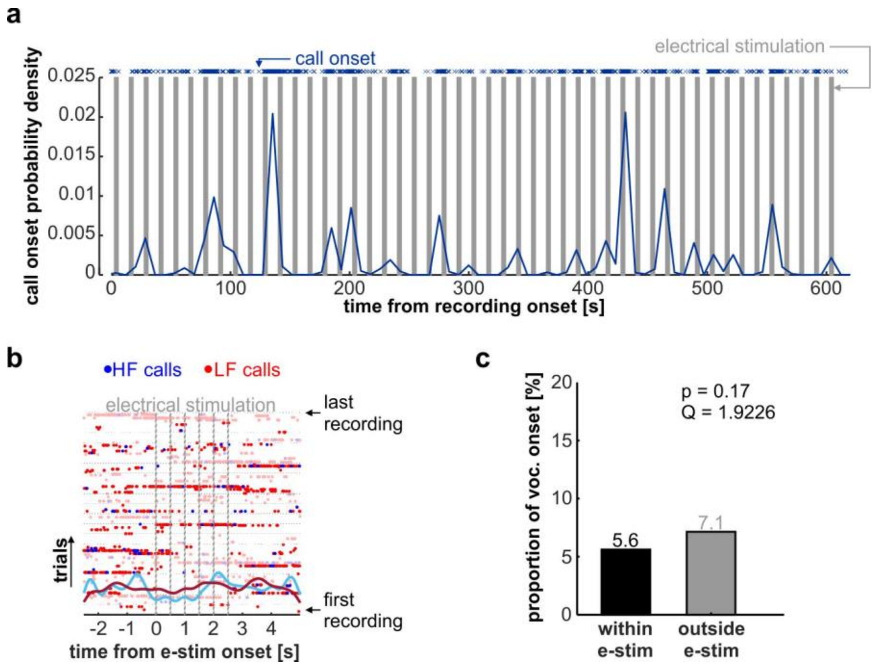


**Fig. S12. Electrical stimulation in FAF does not elicit vocalization.** (**a**) Probability density over the full recording period of call onset, pooled across all recording sessions (n = 3515 calls). Individual call onsets are depicted with an "x" on the top of the plot; electrical stimulation trains (i.e. six pulses) are marked with grey bars. (**b**) Call onset times around periods of electrical stimulation (i.e. during the stimulation train, with pulses illustrated in vertical grey bars) and 2 seconds of pre- and post-time. Calls are shown for each trial of electrical stimulation, sorted from bottom to top, belonging to different recordings (n = 20), also sorted in the same manner. There were 50 trials per recording. Continuous traces illustrate a normalized density function of call onset probability (similar to the one shown in **a**), for low- and high- frequency call (in red and blue, respectively). In panels **a** and **b** no clear effect of electrical FAF stimulation can be observed on the probability of call onset across trials or penetration. (**c**) Proportion of call onset during times of electrical stimulation (black; 5.6%, 55 calls out of 980 trials) vs. proportion of call onset outside times of electrical stimulation (grey; 7.1%, 70 out of 980 trials). We did not see any significant differences in the proportion of call onset across periods ($\chi^2$-test, p = 0.17; Q = 1.9226).

### Supplementary results

### FAF-AC connectivity patterns in pre- and post-vocal periods depend on whether animals vocalize echolocation or communication calls

*Connectivity patterns during pre-vocal periods*

To quantitatively address the differences in preferential information flow shown in **Fig. 2**, we compared connectivity dynamics in the FAF-AC network across vocal conditions (i.e. pre-voc and post-voc; no-voc comparisons shown in **Fig. S5**). The top row of **Fig. S6a** summarizes the outcomes of such comparisons during pre-vocal periods across frequency bands, for the echolocation vs. communication case. Edges in the graphs show significant differences (Wilcoxon rank sum tests, significance when $p < 10^{-4}$) with large effect sizes ($|d| > 0.8$) in the directionality of information flow between two given nodes. Edges were weighted according to the effect size (d) of the corresponding comparisons. Thus, graphs in **Fig. S6a** (top) show that significant differences (with large effect sizes) between the cases of pre-vocal echolocation and pre-vocal communication, in terms of FAF → AC connectivity, occurred only in the $\gamma_2$-band. Within FAF, significant differences in dPTE between were strongest in the $\delta$ and $\alpha$ ranges, although sparse significant differences occurred also in the $\theta$ and $\beta_1$ bands. The strength of preferred FAF → AC directionality of information flow was significantly weaker for pre-vocal echolocation than for no-voc periods in $\delta$ frequencies (**Fig. S5**). However, in the $\gamma_2$ band, FAF → AC connectivity was significantly stronger during pre-vocal echolocation periods. Significant differences in the directionality of information flow between communication and no-voc conditions were rare (**Fig. S5**).

We use the $DI_{net}$ metric (see main text, **Fig.3**) to compare information outflow from each cortical region when animals vocalized either echolocation or communication calls. Significant differences in $DI_{net}$ values across conditions (echolocation vs. communication; **Fig. S6a**, bottom) occurred with large effect sizes ($|d| > 0.8$) only in the $\gamma_2$ band, considering information flowing from the FAF. Specifically, FAF-related $DI_{net}$ in the $\gamma_2$ band was significantly (FDR-corrected Wilcoxon rank sum tests, $p_{corr} < 0.05$) higher when animals vocalized echolocation calls as compared to when communication calls (**Fig. S6a**; $p_{corr} = 4.27 \times 10^{-105}$, $d = 1.83$) or no call whatsoever (**Fig. S5a**; $p_{corr} = 2.24 \times 10^{-84}$, $d = 1.51$). Conversely, $\delta$-band net information outflow was significantly higher during no-voc periods as compared to the pre-vocal echolocation (**Fig. S5a,b**; $p_{corr} = 5.15 \times 10^{-30}$, $d = -0.78$) and the pre-vocal communication (**Fig. S5c**; $p_{corr} = 1.08 \times 10^{-29}$, $d = -0.78$) conditions, although with no large effect sizes in either case.

*Connectivity patterns during post-vocal periods*

There were major differences in connectivity during post-vocal periods between vocalization conditions (**Fig. S6b**). Preferential top-down information flow was significantly lower for echolocation calls than for communication vocalizations in $\delta$ and $\beta_1$ frequencies, but significantly higher in the $\gamma_2$ band (**Fig. S6b**, top; $p < 10^{-4}$, $|d| > 0.8$). Remarkably, post-vocal preferred directionality of information flow in the $\delta$ and $\beta_1$ bands was strongest in the bottom-up direction (AC → FAF) for the echolocation condition. Similar effects were seen when comparing connectivity patterns obtained from post-vocal echolocation and no-voc periods (**Fig. S5c**, top). In other words, the post-vocal echolocation condition exhibited the weakest top-down information transfer and the strongest bottom up-information flow in bands $\delta$ and $\beta_1$. Top-down $\gamma_2$ causal influences remained strongest when animals vocalized an echolocation pulse, as compared to communication call production or no-voc periods. Within-area changes were observed in the $\alpha$-band in FAF, where preferential superficial-to-deep information transfer was significantly higher for echolocation vocalizations (**Fig. S6b**, top), while deep-to-superficial information flow was strongest in post-vocal communication and no-voc related periods (**Fig. S6b, S5**). Finally, significant differences between post-vocal communication and spontaneous activity were limited to $\delta$ frequencies, and strongest for no-voc LFPs.

We compared the net information outflow across conditions in each structure for post-vocal periods (**Fig. S6b**, bottom). In the $\delta$-band, preferred information outflow from the FAF was weakest (with large effect sizes) when animals vocalized echolocation calls (FDR-corrected Wilcoxon rank sum tests; echolocation vs. communication: **Fig. S6b**, $p_{corr} = 1.32 \times 10^{-112}$, $d = -1.66$; echolocation vs. no-voc: **Fig. S5**, $p_{corr} = 1.57 \times 10^{-171}$, $d = -4.27$). A similar effect was observed when comparing communication $DI_{net}$ values with no-voc ones: preferential post-vocal net information outflow from FAF was significantly lower for vocalization-related LFPs (**Fig. S5**, $p_{corr} = 2.58 \times 10^{-125}$, d = -2.22). Similarly, post-vocal $DI_{net}$ values for the $\beta_1$-band in the FAF were significantly stronger during communication than during echolocation production, reaching large effect sizes (**Fig. S6b**, $p_{corr} = 1.71 \times 10^{-38}$, d = -0.85). $\gamma_2$-related net information outflow from FAF was always strongest in the case of echolocation (echolocation vs. communication: **Fig. S6b**, $p_{corr} = 3.8 \times 10^{-144}$, d = 2.69; echolocation vs. no-voc: **Fig. S5**, $p_{corr} = 2.54 \times 10^{-128}$, d = 2.22). In the $\delta$-band, net information outflow from AC was significantly stronger, with large effect sizes, during echolocation production than for post-vocal communication or no-voc periods (echolocation vs. communication: **Fig. S6b** bottom, $p_{corr} = 1.45 \times 10^{-91}$, d = 1.26; echolocation vs. no-voc: **Fig. S5**, $p_{corr} = 2.84 \times 10^{-127}$, d = 1.5). Also, in the $\beta_1$-band, net information outflow from AC was strongest for post-vocal echolocation than communication periods although without large effect sizes (**Fig. S6b**; $p_{corr} = 2.55 \times 10^{-34}$, d = 0.72). Significant changes between echolocation and no-voc cases in the same frequency band did not occur with large effect size (**Fig. S5**; $p_{corr} = 1.84 \times 10^{-12}$, d = 0.37). Differences in other frequency bands, or other across-condition comparisons (e.g. communication vs. no-voc), were either not reflected in the differential connectivity graphs, or did not have large effect sizes.

Altogether, these results indicate that pre-vocal and post-vocal directional information flow in the FAF-AC network occurs mostly in low and high-frequency bands. The patterns and strength of preferred directionality depend on whether a vocalization is produced and on the type of vocal output. When animals produced echolocation calls, post-vocal bottom-up influences dominated in $\delta$ frequencies, while top-down influences weakened in post-vocal periods compared to spontaneous activity. These results could reflect both a waning of top-down control from the FAF, and an increase in bottom-up transfer in $\delta$ and $\beta_1$ frequencies. These two possible explanations are not mutually exclusive.

**<u>Passive listening of high- or low-frequency natural sounds does not explain information flow patterns of active vocalization</u>**

Recordings for this study were made mostly from an area of the *C. perspicillata*'s AC whose neurons are specialized for processing echolocation sounds. Therefore, it is sensible to assume that the reversal of preferred directionality of information flow from pre- to post-vocal periods during vocalization could be attributed to strong acoustic feedback originating from an echolocation call, interacting with the tuning of the cortical areas recorded. In the following, we present evidence demonstrating that mere auditory feedback is not sufficient to explain our main results.

In a first step, we quantified frequency tuning in the AC (and FAF, see Methods) and observed the tuning of recorded LFPs did not favour the frequency range of echolocation calls (i.e. > 60 kHz), as it peaked at 20-40 kHz for most recording sites (**Fig. S7** shows LFP frequency tuning curves measured with 75 dB SPL, 10 ms tones, across penetrations). Thus, LFP responses in the AC, at least based on frequency tuning alone, would not elicit on average a stronger response and therefore a stronger bottom-up transfer towards the FAF. Note that many recordings were responsive at high frequencies, often exhibiting double-peaked tuning curves (e.g. red trace in **Fig. S7a**). This type of tuning is common in the auditory systems of *C. perspicillata* and other bats (Kanwal et al., 1999; Lopez-Jury et al., 2021; Radtke-Schuller and Schuller, 1995), potentially facilitating neurons to respond to both echolocation and communication sounds. In the FAF, we did not observe clear frequency tuning based on LFPs.

In a second step, we quantified information transfer dynamics in the FAF-AC network in response to acoustic stimulation. Sounds were a high-frequency frequency-modulated sweep ("HF-FM"; intended to mimic an echolocation pulse) and a natural distress syllable ("distress"; as a representative of a communication utterance). These sounds are illustrated in **Fig. S7b**, together with simultaneous cortical responses to each from FAF and AC in **Fig. S7c**. When considering LFPs taken in the period from 0-500 ms after stimulus onset, we observed that information flowed predominantly in the FAF → AC direction for bands δ, θ, and $\gamma_2$ regardless of the sound considered (**Fig S7d**). The patterns of information flow were very similar across the types of acoustic stimuli used, although with some significant differences in the α and $\gamma_2$ bands (quantified in **Fig. S8**). Overall, preferential information flow dynamics in these and other frequencies were reminiscent of those observed for no-voc periods, as confirmed by scarce differences between information flow patterns associated to passive stimulation and spontaneous activity (**Fig. S8**).

Preferential information flow patterns associated to post-vocal echolocation periods were significantly different than those reported for the passive listening of HF-FM sounds (**Fig. S9a**). Specifically, for δ frequencies, information flow in the FAF → AC direction was stronger in the passive listening condition, but stronger in the AC → FAF direction for post-vocal echolocation periods. Differences were significant with large effect sizes also when considering the net information outflow across structures in this band ($p \leq 1.12 \times 10^{-127}$, $|d| \geq 1.5$; **Fig S9a**, bottom). In the θ band, predominant information flow was strongest in the FAF→AC direction when animals listened to the HF-FM stimulus ($p = 3.45 \times 10^{-58}$, $d = 1.17$). However, information transfer in the FAF→AC direction was strongest for post-vocal echolocation periods in the $\gamma_2$ band (comparison of $DI_{net}$ values: $p = 6.98 \times 10^{-100}$, $d = -1.72$). Passive listening of distress syllables yielded stronger information flow in the FAF → AC direction when compared to post-vocal communication periods in the δ and θ frequency bands (comparison of $DI_{net}$ values: $p \leq 5.29 \times 10^{-37}$, $d \geq 0.88$; **Fig. S9b**, bottom). For $\beta_1$ frequencies, information flowed more strongly in the AC→FAF direction considering post-vocal communication periods, but the differences in $DI_{net}$ values did not occur with large effect sizes ($p = 1.83 \times 10^{-18}$, $d = 0.52$).

Altogether, these results demonstrate that acoustic input (as may occur from feedback after call utterance) does not account for the reversal of information transfer in δ frequencies when animals produce echolocation calls. Our results also provide evidence for a highly dynamic network, in which information reverses in different manners during vocalization and passive listening.

### Information transfer patterns related to HF-communication calling differ to those associated with echolocation

The passive listening of high frequency acoustic stimuli does not explain the information transfer patterns associated to echolocation production (**Fig. S7-9**), but it is possible that the utterance of high frequency sounds in general (irrespective of whether echolocation or communication) suffices to account for such dynamics. To explore this possibility, we quantified information transfer in the FAF-AC network when animals produced HF-communication sounds.

Post-vocal information transfer dynamics associated to the production of HF-communication vocalizations are shown in **Fig. S10a**. Mostly for frequency bands θ and α (and more weakly, $\beta_1$), it is apparent that information flows predominantly in the AC → FAF direction. When compared to pre-vocal periods (**Fig. S10b**, top), we observed pre-vocal to post-vocal switch in directionality across FAF and AC for θ-band LFPs, but not for δ-band ones (i.e. the band where the reversal of information flow occurred when producing echolocation pulses). The described effect in θ was complemented by differences in $DI_{net}$ values ($p = 1.25x10^{-45}$, $d = -0.99$; **Fig. S10b**, bottom). Indeed, a comparison between patterns of predominant information transfer associated to HF-communication and echolocation production (**Fig. S10c**), revealed that information flowed more strongly for echolocation production than for HF-communication utterance in δ frequencies (also complemented by $DI_{net}$ values: $p = 1.66x10^{-70}$, $d = -1.12$). Moreover, small differences were found in the case of AC → FAF dPTE values in the θ band, indicating that bottom-up information transfer for post-vocal echolocation and HF-communication periods were not considerably different from one another. Additional differences between the conditions of echolocation and HF-communication production were observed in other frequency bands (see α, $\beta_1$, and $\gamma_2$ in **Fig. S10c**), although for these frequencies there were no significant differences between pre- and post-vocal periods when animals vocalized HF-communication sounds (**Fig. S10b**). Taken together, these results indicate that the information transfer dynamics in the FAF-AC circuit observed when animals vocalize echolocation pulses are not accounted for solely by the frequency content of the calls produced by the bats.

Kanwal, J.S., Fitzpatrick, D.C., and Suga, N. (1999). Facilitatory and inhibitory frequency tuning of combination-sensitive neurons in the primary auditory cortex of mustached bats. J Neurophysiol *82*, 2327-2345.
Lopez-Jury, L., Garcia-Rosales, F., Gonzalez-Palomares, E., Kossl, M., and Hechavarria, J.C. (2021). Acoustic Context Modulates Natural Sound Discrimination in Auditory Cortex through Frequency-Specific Adaptation. J Neurosci *41*, 10261-10277.
Radtke-Schuller, S., and Schuller, G. (1995). Auditory cortex of the rufous horseshoe bat: 1. Physiological response properties to acoustic stimuli and vocalizations and the topographical distribution of neurons. Eur J Neurosci *7*, 570-591.
